# Distinct ipRGC subpopulations mediate light’s acute and circadian effects on body temperature and sleep

**DOI:** 10.1101/496323

**Authors:** Alan C. Rupp, Michelle Ren, Cara M. Altimus, Diego C. Fernandez, Melissa Richardson, Fred Turek, Samer Hattar, Tiffany Schmidt

**Affiliations:** Department of Biology, Johns Hopkins University, Baltimore, MD, USA.; Department of Neurobiology, Northwestern University, Evanston, IL, USA; Department of Neuroscience, Johns Hopkins University, Baltimore, MD, USA.

## Abstract

The light environment greatly impacts human alertness, mood, and cognition by acute regulation of physiology and indirect alignment of circadian rhythms. Both processes require the melanopsin-expressing intrinsically photosensitive retinal ganglion cells (ipRGCs), but the relevant downstream brain areas remain elusive. ipRGCs project widely in the brain, including to the central circadian pacemaker, the suprachiasmatic nucleus (SCN). Here we show that body temperature and sleep responses to light are absent after genetic ablation of all ipRGCs except a subpopulation that projects to the SCN. Furthermore, by chemogenetic activation of the ipRGCs that avoid the SCN, we show that these cells are sufficient for acute changes in body temperature. Our results challenge the idea that the SCN is a major relay for the acute effects of light on non-image forming behaviors and identify the sensory cells that initiate light’s profound effects on body temperature and sleep.

## Introduction

Many essential functions are influenced by light both indirectly through alignment of circadian rhythms (photoentrainment) and acutely by a direct mechanism (sometimes referred to as ‘masking’) (Mrosovsky et al., 1999; Altimus et al., 2008; Lupi et al., 2008; Tsai et al., 2009; Legates et al., 2012). Dysregulation of the circadian system by abnormal lighting conditions has many negative consequences, which has motivated decades of work to identify the mechanisms of circadian photoentrainment (Golombek and Rosenstein, 2010). In contrast, it has only recently become apparent that light exposure can also acutely influence human alertness, cognition, and physiology (Chellappa et al., 2011). As a result, there is a developing awareness of light quality in everyday life (Lucas et al., 2014). It is therefore essential to human health and society to elucidate the circuitry and coding mechanisms underlying light’s acute effects.

Intriguingly, a single population of retinal projections neurons—intrinsically photosensitive retinal ganglion cells (ipRGCs)—have been implicated in the circadian and acute effects of light on many functions, including activity, sleep, and mood (Göz et al., 2008; Güler et al., 2008; Hatori et al., 2008; Legates et al., 2012; Fernandez et al., 2018). ipRGCs integrate light information from rods, cones, and their endogenous melanopsin phototransduction cascade (Schmidt et al., 2011), and relay that light information to over a dozen central targets (Hattar et al., 2006; Ecker et al., 2010). However, the circuit mechanisms mediating ipRGC-dependent functions are largely unknown.

One notable exception is the control of circadian phototentrainment. It is accepted that ipRGCs mediate photoentrainment by direct innervation of the master circadian pacemaker, the suprachiasmatic nucleus (SCN) of the hypothalamus (Göz et al., 2008; Güler et al., 2008; Hatori et al., 2008; Jones et al., 2015). This is supported by studies demonstrating that genetic ablation of ipRGCs results in mice with normal circadian rhythms that ‘free-run’ with their endogenous rhythm, independent of the light/dark cycle (Göz et al., 2008; Güler et al., 2008; Hatori et al., 2008). Further, mice with genetic ablation of all ipRGCs except those that project to the SCN and intergeniculate leaflet (IGL) display normal circadian photoentrainment (Chen et al., 2011), suggesting that ipRGC projections to the SCN/IGL are sufficient for photoentrainment.

In comparison, the mechanisms by which ipRGCs mediate acute light responses remains largely a mystery. Genetic ablation of ipRGCs or their melanopsin phototransduction cascade blocks or attenuates the acute effects of light on sleep (Altimus et al., 2008; Lupi et al., 2008; Tsai et al., 2009), wheel-running activity (Mrosovsky and Hattar, 2003; Güler et al., 2008), and mood (Legates et al., 2012; Fernandez et al., 2018). This dual role of ipRGCs in circadian and acute light responses suggests they may share a common circuit mechanism. However, whether the circuit basis for ipRGCs in the acute effects of light and circadian functions is through common or divergent pathways has yet to be determined. ipRGCs project broadly in the brain beyond the SCN (Hattar et al., 2002, 2006; Gooley et al., 2003; Baver et al., 2008). Additionally, ipRGCs are comprised of multiple subpopulations with distinct genetic, morphological, and electrophysiological signatures (Baver et al., 2008; Schmidt and Kofuji, 2009; Ecker et al., 2010; Schmidt et al., 2011) and distinct functions (Chen et al., 2011; Schmidt et al., 2014). Though there are rare exceptions (Chen et al., 2011; Schmidt et al., 2014), the unique roles played by each ipRGC subsystem remain largely unknown.

It is currently unknown whether distinct ipRGC subpopulations mediate both the acute and circadian effects of light, and two major possibilities exist for how this occurs: (1) ipRGCs mediate both acute and circadian light responses through their innervation of the SCN or (2) ipRGCs mediate circadian photoentrainment through the SCN, but send collateral projections elsewhere in the brain to mediate acute light responses. To date, the predominant understanding has centered on a role for the SCN in both acute and circadian responses to light (Muindi et al., 2014; Morin, 2015). However, this model has been controversial due to complications associated with SCN lesions (Redlin and Mrosovsky, 1999) and alternative models proposing a role for direct ipRGC input to other central targets (Redlin and Mrosovsky, 1999; Lupi et al., 2008; Tsai et al., 2009; Hubbard et al., 2013; Muindi et al., 2014). Here, we sought to address the question of how environmental light information—through ipRGCs—mediates both the circadian and acute regulation of physiology. To do so, we investigated the ipRGC subpopulations and coding mechanisms that mediate body temperature and sleep regulation by light. We find that a molecularly distinct subset of ipRGCs is required for the acute, but not circadian, effects of light on thermoregulation and sleep. These findings suggest that, contrary to expectations, functional input to the SCN is not sufficient to drive the acute effects of light on these behaviors. These findings provide new insight into the circuits through which light regulates behavior and physiology.

## Results

### Brn3b-positive ipRGCs are required for light’s acute effects on thermoregulation

To identify mechanisms of acute thermoregulation, we maintained mice on a 12-hr/12-hr light/dark cycle and then presented a 3-hr light pulse two hours into the night (Zeitgeber time 14, ZT14) while measuring core body temperature (Fig. 1A). The nocturnal light pulse paradigm is well-established for studying acute regulation of sleep and wheel-running activity (Mrosovsky et al., 1999; Mrosovsky and Hattar, 2003; Altimus et al., 2008; Lupi et al., 2008). We focused first on body temperature because of its critical role in cognition and alertness (Wright et al., 2002; Darwent et al., 2010), sleep induction and quality (Kräuchi et al., 1999), metabolic control (Kooijman et al., 2015), and circadian resetting (Buhr et al., 2010).

**Figure 1:**
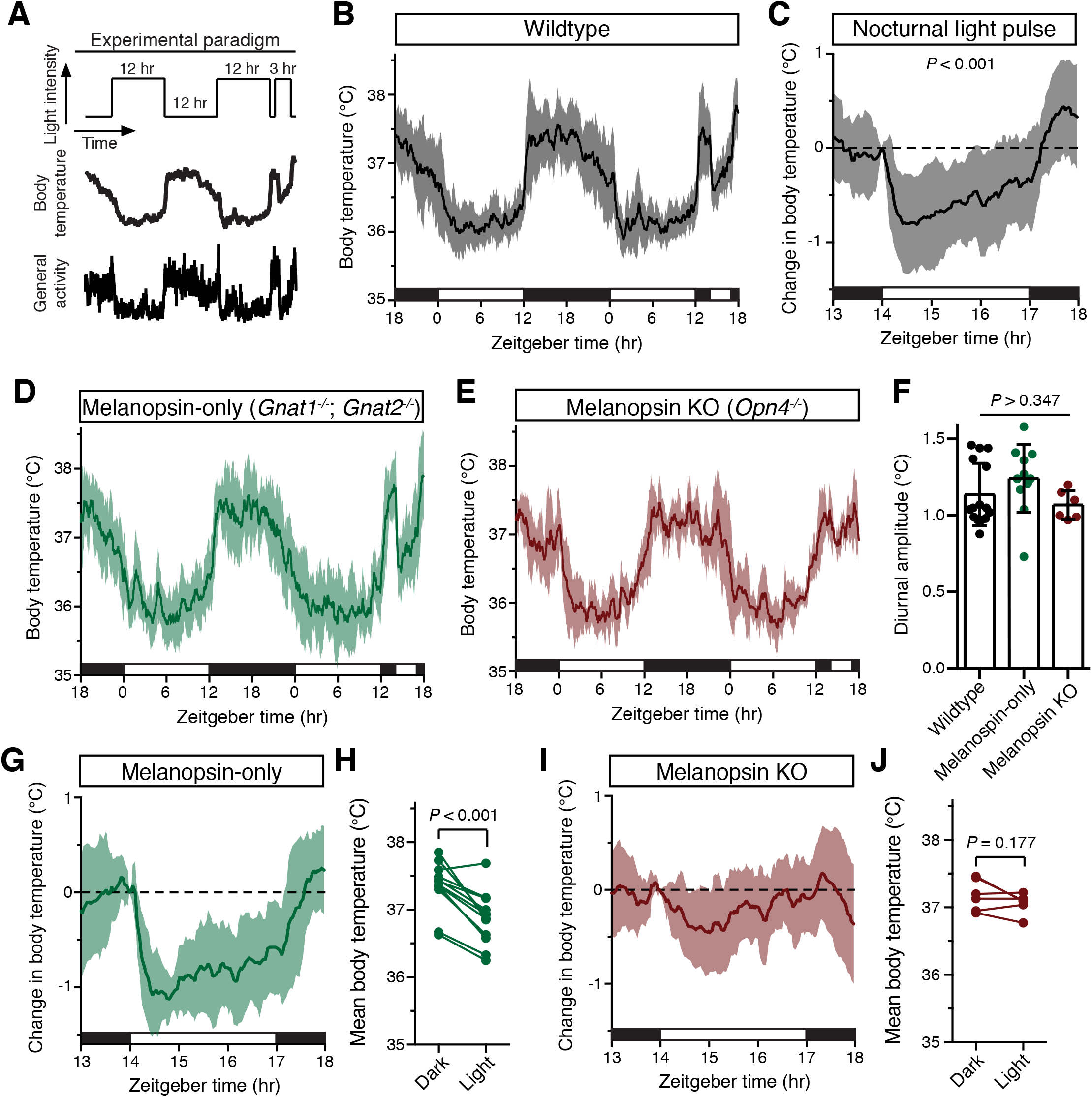
Melanopsin mediates the acute effects of light on body temperature. (**A**) Paradigm to measure body temperature continuously in a 12:12 light dark cycle with a 3-hour light pulse at ZT14. (**B**) 48 hours of continuous body temperature monitoring in wildtype male mice (n = 13) (**C**) Body temperature in WT during light pulse, compared to baseline (ZT14). *P* < 0.001, paired t-test of mean temperature compared to previous night. (**E**) Melanopsin-only mice (*Gnat1*^-/-^; *Gnat2*^-/-^, n = 11) and (**E**) melanopsin knockout (*Opn4*^-/-^, n = 6) 48-hour diurnal body temperature. (**F**) Diurnal body temperature amplitude in the three groups. *P* > 0.347 for effect of group by one-way ANOVA. (**G**) Body temperature in melanopsin-only during light pulse, relative to baseline (ZT14). (**H**) Paired comparison of body temperature during light pulse compared to previous night. *P* < 0.001 by paired t-test. (**I**) Body temperature in melanopsin knockout during light pulse, relative to baseline (ZT14). (**J**) Paired comparison of body temperature during light pulse compared to previous night.

Body temperature photoentrains to the light/dark cycle with peaks during the night and troughs during the day (Fig. 1B). Both rodents and humans utilize ocular light detection to acutely adjust body temperature in response to a nocturnal light pulse (Dijk et al., 1991; Cajochen et al., 2005), though how this body temperature change is initiated by the retina and relayed to the brain is unknown. When we presented wildtype mice with a nocturnal light pulse, we observed a decrease in both body temperature and general activity compared to the previous night (Fig. 1C). The decrease in body temperature and activity was sustained for the entire 3-hr stimulus, with moderate rundown (Fig. 1C).

We observed that acute body temperature regulation only occurred at relatively bright light intensities (>100 lux) (Fig. S1). This, in combination with previous reports that body temperature regulation is most sensitive to short-wavelength light (Cajochen et al., 2005), suggested that it might be mediated by the insensitive and blue-shifted melanopsin phototransduction (Lucas et al., 2001; Do et al., 2009). To test this, we measured body temperature in mice lacking either functional rods and cones (melanopsin-only: *Gnat1^-/-^*; *Gnat2^-/-^*) or lacking melanopsin (melanopsin KO: *Opn4^-/-^*). Both genotypes photoentrained their body temperature (Fig. 1 D,E), with an amplitude indistinguishable from wildtype (Fig. 1F). However, we found that acute body temperature decrease to a nocturnal light pulse was present in melanopsin-only mice (*Gnat1*^-/-^; *Gnat2*^-/-^) (Fig. 1G,H), but absent from melanopsin knockout mice (*Opn4^-/-^*) (Fig. 1I,J). This indicates that melanopsin is critical for light’s ability to drive acute body temperature decreases, as it is for acute sleep induction (Altimus et al., 2008; Lupi et al., 2008; Tsai et al., 2009). These results suggest that ipRGCs are the only retinal cells that are necessary and sufficient for acute thermoregulation by light.

ipRGCs comprise multiple subtypes with distinct gene expression profiles, light responses, and central projections (Schmidt et al., 2011), prompting us to ask which subtypes mediate acute thermoregulation. Brn3b(+) ipRGCs project to many structures including the olivary pretectal nucleus (OPN) and dorsal lateral geniculate nucleus (dLGN), but largely avoid the SCN (Chen et al., 2011). In contrast, Brn3b(-) ipRGCs project extensively to the SCN and intergeniculate leaflet (IGL), while avoiding the OPN and dLGN (Chen et al., 2011). Ablation of Brn3b(+) ipRGCs using melanopsin-Cre and a Cre-dependent diphtheria toxin knocked into the *Brn3b* locus (Brn3b-DTA: *Opn4*^*Cre*/+^;*Brn3b*^*zDTA*/+^) removes virtually all ipRGC input to brain areas aside from the SCN and IGL, and these mice retain circadian photoentrainment of wheel-running activity (Chen et al., 2011).

When we measured body temperature in Brn3b-DTA mice, we found that their body temperature was photoentrained with a similar amplitude to controls (Fig. 2A-C). However, despite the presence of melanopsin in Brn3b-DTA mice (*Opn4*^*Cre*/+^;*Brn3b*^*zDTA*/+^), they did not acutely decrease body temperature in response to a nocturnal light pulse (Fig. 2F,G). Importantly, melanopsin heterozygous littermate controls (*Opn4*^*Cre*/+^) displayed normal acute thermoregulation by light (Fig. 2D,E), indicating that halving melanopsin gene dosage is not the cause of the impaired body temperature decrease in Brn3b-DTA mice. These results demonstrate that Brn3b(+) ipRGCs are required for acute thermoregulation regulation by light but not photoentrainment of body temperature and reveal that light information to the SCN is sufficient for circadian photoentrainment of body temperature, but not its acute regulation.

**Figure 2:**
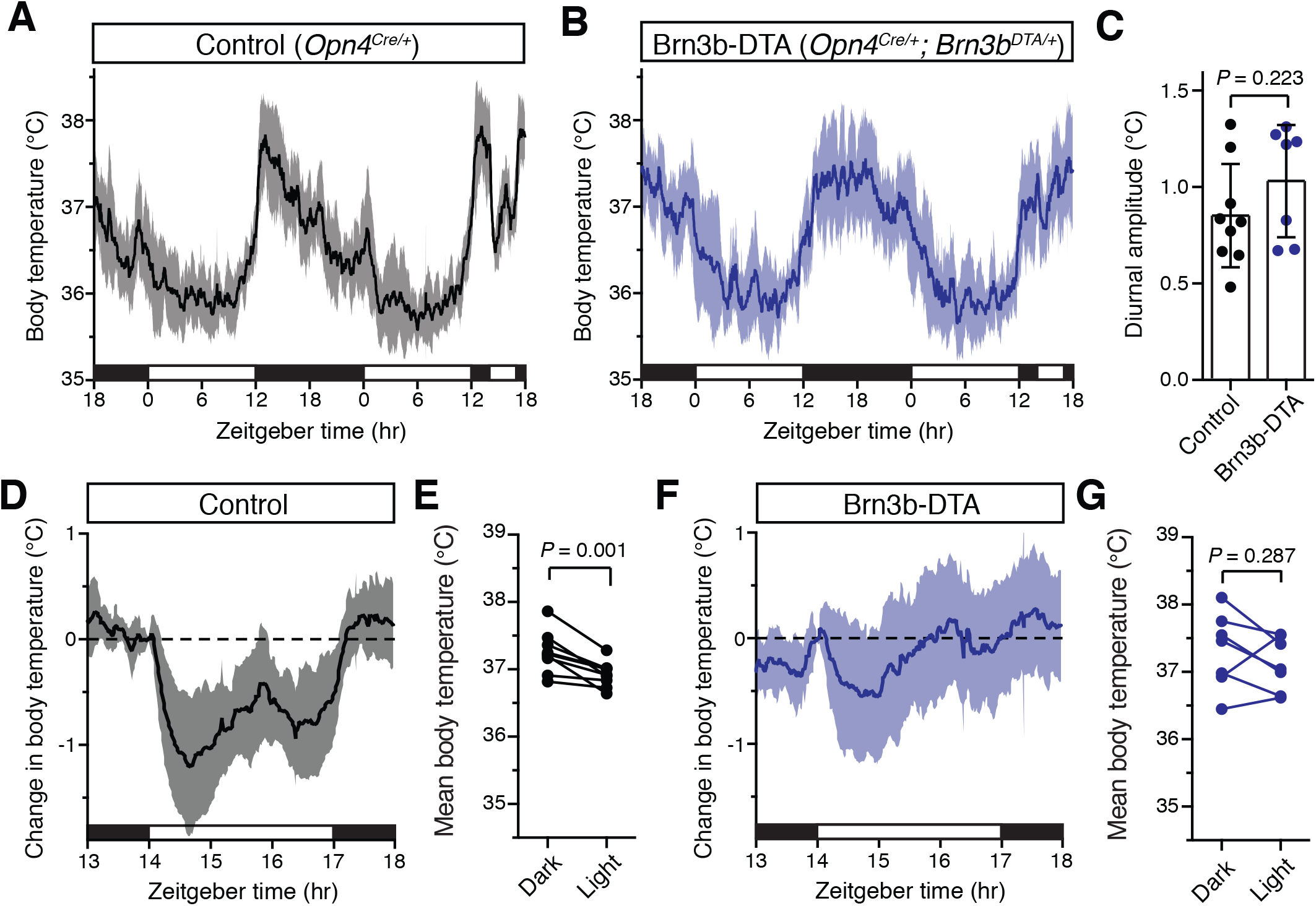
Brn3b-negative ipRGCs are insufficient for acute body temperature regulation via the SCN. (**A**) Diurnal body temperature in control (*Opn4*^*Cre*/+^, n = 9) and (**B**) Brn3b-DTA (*Opn4*^*Cre*/+^;*Brn3b*^*DTA*/+^, n = 7). (**C**) Diurnal body temperature amplitude in the two groups. *P* = 0.223 by t-test. (**D**) Body temperature in control during light pulse, relative to baseline (ZT14). (**E**) Paired comparison of body temperature during light pulse compared to previous night. *P* = 0.001 by paired t-test. (**F**) Body temperature in Brn3b-DTA during light pulse, relative to baseline (ZT14). (**G**) Paired comparison of body temperature during light pulse compared to previous night. *P* = 0.287 by paired t-test.

### Brn3b-positive ipRGCs are sufficient for acute thermoregulation

Our data thus far suggest that there are two functionally distinct populations of ipRGCs that regulate the same physiological function: (1) Brn3b(-) ipRGCs that project to the SCN to mediate circadian photoentrainment of body temperature and (2) Brn3b(+) ipRGCs that project elsewhere in the brain to mediate acute thermoregulation. To test if Brn3b(+) ipRGCs are sufficient for acute thermoregulation, we expressed a chemogenetic activator in Brn3b(+) RGCs (Fig. 3A, *Brn3b*^*Cre*/^+ with intravitreal AAV2-hSyn-DIO-hM3Dq-mCherry, we refer to these mice as Brn3b-hM3Dq. This technique allows us to acutely activate the Brn3b(+) RGCs with the DREADD agonist clozapine N-oxide (CNO) (Armbruster et al., 2007). We found that after intravitreal viral delivery, many RGCs were infected, including melanopsin-expressing ipRGCs (Fig. 3A).

**Figure 3:**
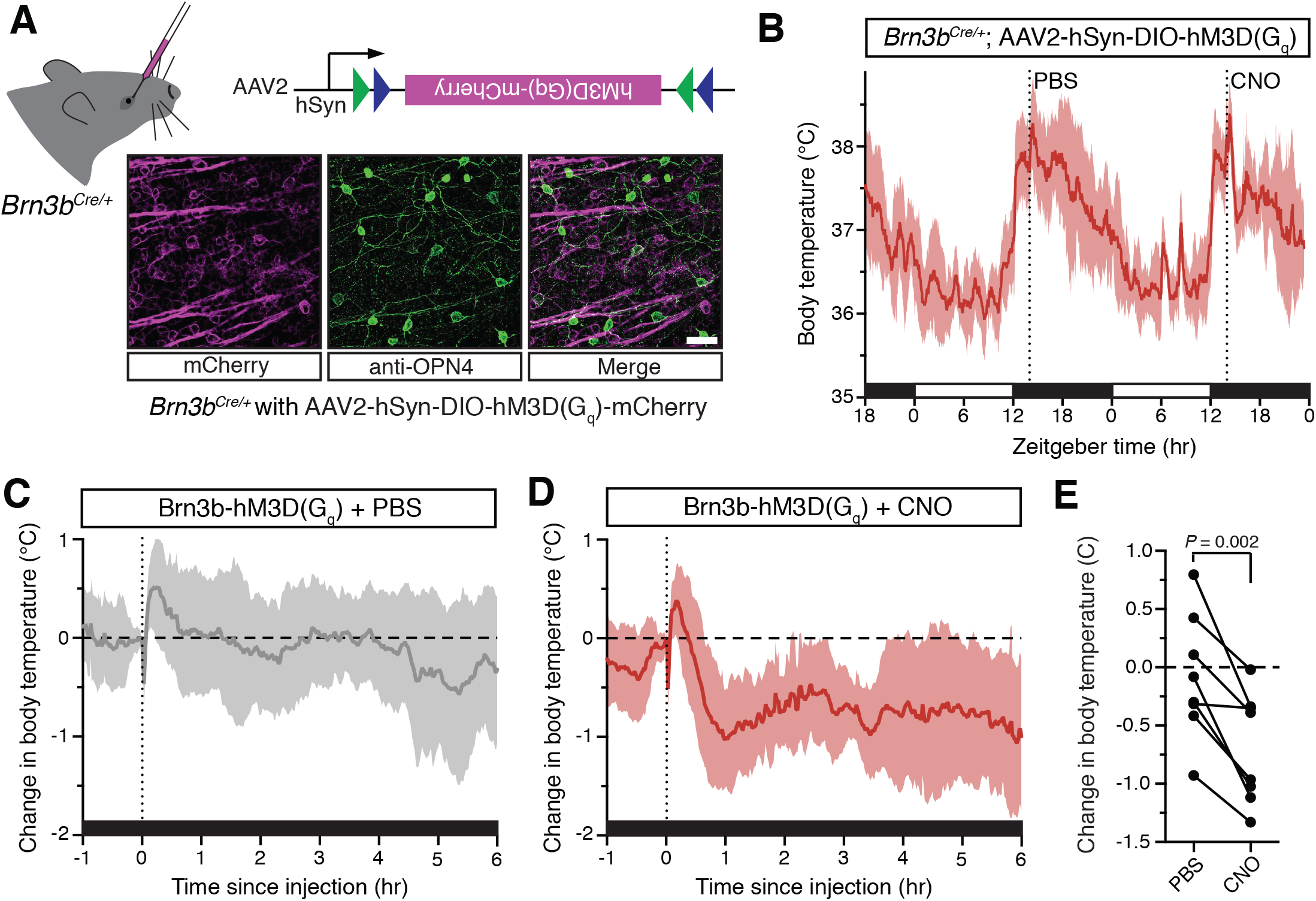
Activation of Brn3b-positive RGCs is sufficient to drive sustained body temperature decreases. (**A**) Diagram of intravitreal delivery of AAV2-hSyn-DIO-hM3Dq-mCherry to *Brn3b*^*Cre*/+^ mice, and confirmation of infection of ipRGCs. (**B**) 54-hr continuous diurnal body temperature recordings in Brn3b-hM3Dq mice, with injections of PBS then CNO on consecutive nights at ZT14. (**C**) Change in body temperature after PBS injection, relative to baseline (time of injection). (**D**) Change in body temperature after CNO injection, relative to baseline (time of injection). (**E**) Paired comparison of the change in body temperature with either PBS or CNO injection. *P* = 0.002 by paired t-test.

Importantly, Brn3b-hM3Dq mice photoentrained to a normal light/dark cycle (Fig. 3B). Following CNO administration at ZT14 to depolarize the Brn3b(+) RGCs, we observed a robust decrease in body temperature that lasted at least 6 hours (Fig. 3D). Importantly, PBS administration in Brn3b-hM3Dq mice (Fig. 3C) and nocturnal CNO administration in wildtype control mice (Fig. S2) had no measurable effect on body temperature. Together, these results demonstrate that Brn3b(+) ipRGCs mediate the acute effects of light on body temperature though extra-SCN projection(s), while Brn3b(-) ipRGCs mediate circadian photoentrainment by projections to the SCN and/or IGL.

### Brn3b-positive ipRGCs are required for light’s acute effects on sleep

We next examined the contribution of Brn3b(+) and Brn3b(-) ipRGCs to sleep. To do this, we used EEG and EMG recordings to compare the sleep behavior of Control (*Opn4*^*Cre*/+^) and Brn3b-DTA mice. We first analyzed the daily sleep patterns and proportion of rapid eye movement (REM) and non-REM (NREM) sleep in Control and littermate Brn3b-DTA animals. We found that Brn3b-DTA mice show normal photoentrainment of sleep and similar percent time of sleep across the 24 hour day, with only one 30 minute bin at ZT12 (light offset) showing a significant difference between Control and Brn3b-DTA animals (Fig. 4A,B). This is consistent with previous reports of normal circadian photoentrainment of daily activity rhythms in Brn3b-DTA mice (Chen et al., 2011). Control and Brn3b-DTA mice also showed similar total percent time awake or asleep across an entire day (Fig. 4C), though Brn3b-DTA mice showed a small, but significant, increase in the proportion of total sleep that was classified as NREM and decrease in the proportion of total sleep that was classified as REM (Fig. S3A).

**Figure 4:**
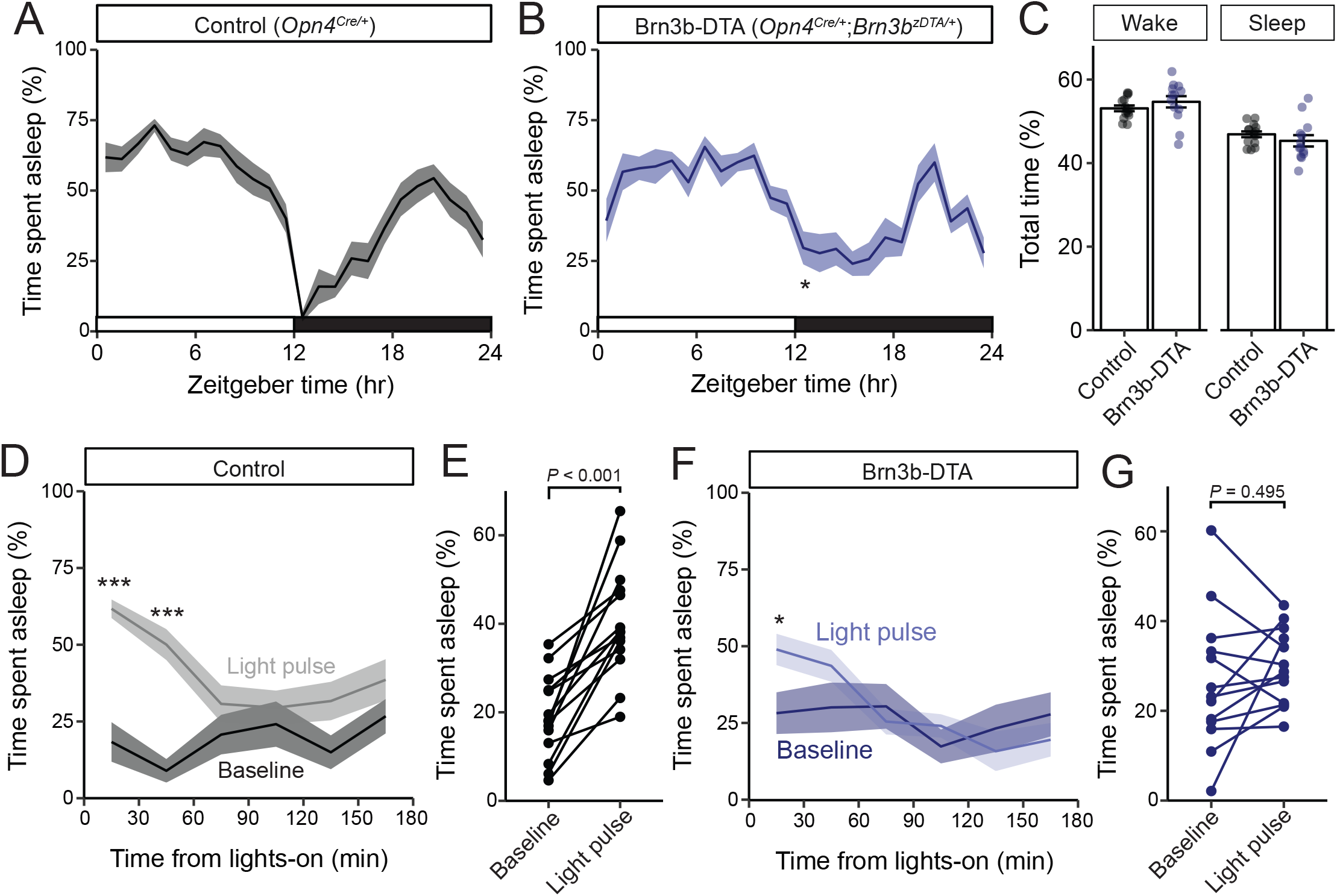
Brn3b-positive M1 ipRGCs are not required for circadian photoentrainment of sleep, but are required for its acute induction by light. (A-C) Percent time spent asleep in 1-hr bins across the 24-hr day for (A) Control (black) mice (n = 14) and (B) Brn3b-DTA (blue) mice (n = 13) lacking Brn3b-positive ipRGCs. Both lines showed normal photoentrainment of sleep, with no main effect of genotype compared to Control by repeated-measures two-way ANOVA (F (1, 25) = 1.108, *P* = 0.303). Brn3b-DTA mice showed a significant reduction in sleep only at lights off (ZT 12) by Sidak’s multiple comparisons test (*P* = 0.029). (C) Percent time spent awake and asleep in Control (black) and Brn3b-DTA mice (blue). No differences were observed between genotypes by t-test (*P* = 0.316). (D-G) Percent time spent asleep for (D) Control mice (black) and (F) Brn3b-DTA mice (blue) at baseline (dark line) and during the three hour light pulse (light line). Significant difference from baseline determined by repeated measures two-way ANOVA. Significant effect of treatment for Controls (F (1, 13) = 38.09, P < 0.001), but not for Brn3b-DTA (F (1, 12) = 0.8496, *P* = 0.375). (E) Control mice show significantly more sleep and less wake during a light pulse (paired t-test) while (G) Brn3b-DTA mice showed no change in percent sleep or wake during the same period. Data is mean for ZT14-17.

We hypothesized that this small difference in sleep at lights-off in Brn3b-DTA mice could be due to a defect in their acute response to light for sleep modulation. To test this, we subjected mice to a 3-hr light pulse from ZT14-17 (Altimus et al. 2008), when the homeostatic drive for sleep is low and Control and Brn3b-DTA animals display similar amounts of sleep (Fig. 4A,B). We found that in Control mice, a light pulse decreased time awake and increased time asleep relative to baseline (previous day) (Fig. 4C,D), while in Brn3b-DTA mice a light pulse caused no change in total percent time asleep or awake (Figure 4F,G), but moderately increased sleep in the first 30-min bin (Fig. 4F). Neither Control nor Brn3b-DTA animals showed any change in proportion of non-REM or REM sleep in response to the light pulse (Fig. S3B,C). These data show that Brn3b(+) ipRGCs are necessary for the acute light induction of sleep. Consistent with our body temperature data, although Brn3b-DTA mice have apparently normal input to the SCN and show normal circadian photoentrainment of wheel-running activity (Chen et al., 2011), body temperature (Fig. 2), and sleep (Fig. 4), this ipRGC innervation of the SCN is not sufficient to drive the normal light induction of sleep. These disruptions in light’s acute effects on thermoregulation and sleep are circuit specific effects because Brn3b-DTA mice showed robust inhibition of wheel running behavior to a 3-hr light pulse delivered from ZT14-17 (Fig. S4).

## Discussion

We show here that for the same physiological outcome, the acute effects of light are relayed through distinct circuitry from that of circadian photoentrainment, despite both processes requiring ipRGCs. Our results suggest that for thermoregulation and sleep, ipRGCs can be genetically and functionally segregated into Brn3b(+) ‘acute’ cells, and Brn3b(-) ‘circadian’ cells. Because Brn3b(+) cells largely avoid the SCN, and Brn3b(-) cells preferentially target the SCN, our results discount a critical role for the SCN in acute light responses in these two behaviors, and instead implicate direct ipRGC projections to other brain areas (Gooley et al., 2003; Hattar et al., 2006). Surprisingly, Brn3b(-) cells are sufficient to drive the acute and circadian effects of light on wheel running activity, demonstrating further divergence in the circuits mediating the acute effects of light on behavior, and suggesting that, unlike for thermoregulation and sleep, acute and circadian regulation of activity is driven via the SCN.

The specific Brnb(+) ipRGC subtypes that mediate the light’s acute effects on body temperature and sleep remain a mystery. A majority of all known ipRGC subtypes (M1-M5) are lost in Brn3b-DTA mice (Chen et al., 2011), with the exception of a subset of ~200 M1 ipRGCs. In agreement with this, ipRGC projections to all minor hypothalamic targets are lost in Brn3b-DTA mice, while innervation of the SCN and part of the IGL remains intact (Chen et al. 2011, Li and Schmidt, 2018). This suggests that all non-M1 subtypes and a portion of M1 ipRGCs are Brn3b(+). Each subtype has a distinct reliance on melanopsin versus rod/cone phototrasnsduction for light detection (Schmidt and Kofuji, 2009). The necessity and sufficiency of melanopsin in mediating acute effects of light on body temperature (Fig. 1) and sleep (Altimus et al., 2008; Lupi et al., 2008; Tsai et al., 2009) suggests that a subtype with strong melanopsin, but weak rod/cone photodetection is responsible - possibly either M1 or M2 cells. However, experiments to tease this apart will require novel methods to specifically manipulate ipRGC subtypes that are currently unavailable.

The brain areas that mediate the acute effects of light on physiology are essentially unknown. There are many candidate areas that both receive direct ipRGC innervation and have been documented to be involved in light’s acute effects on physiology, including the preoptic areas (Muindi et al. 2014), the ventral subparaventricular zone (Kramer et al., 2001), and the pretectum/superior colliculus (Miller et al., 1998). Aside from the SCN, ipRGC projections to the median (MPO) and ventrolateral preoptic (VLPO) areas have been the most widely supported. The preoptic areas are involved in sleep and body temperature regulation (Szymusiak and McGinty, 2008; Nakamura, 2011) and are activated by an acute light pulse (Lupi et al., 2008; Tsai et al., 2009). In support of our behavioral findings, ipRGC projections to each of these areas is lost in Brn3b-DTA animals (Li and Schmidt, 2018). However, ipRGC projections to these areas are sparse (Gooley et al., 2003; Hattar et al., 2006), suggesting their activation by light may be indirect.

In contrast, the superior colliculus (SC) and pretectum receive robust innervation from ipRGCs (Hattar et al., 2002, 2006; Gooley et al., 2003; Ecker et al., 2010), their lesioning blocks light’s acute effects on sleep (Miller et al., 1998), and melanopsin knockout mice lose light-induced cFOS expression in the SC (Lupi et al., 2008). However, the SC and pretectum receive robust innervation from many conventional RGCs, making the requirement for melanopsin and ipRGCs in acute sleep and body temperature regulation difficult to reconcile. It is also possible (and perhaps probable), that multiple ipRGC target regions are involved, with distinct areas mediating distinct physiological responses. Future studies silencing each retinorecipient target while activating Brn3b(+) ipRGCs will be necessary to tease apart the downstream circuits mediating light’s acute effects on physiology.

Alternatively, it remains possible that direct ipRGC control of body temperature is the primary and critical step for many acute responses to light that are mediated by ipRGCs. In support of this possibility, changes in body temperature and heat loss can directly influence sleep induction (Kräuchi et al., 1999). This change in sleep is in turn presumably causative of at least some of light’s effects on wheel-running and general activity (Mrosovsky et al., 1999). Further, core body temperature can acutely regulate cognition and alertness (Wright et al., 2002; Darwent et al., 2010). It is therefore possible that ipRGCs can have widespread influence on an animal’s basic physiology and cognitive function simply by regulating body temperature.

Together, our identification of the photopigment and the retinal circuits mediating acute body temperature and sleep induction will facilitate better methods to promote or avoid human alertness and cognition at appropriate times of day (Chellappa et al., 2011). Our results support many recent efforts to capitalize on the specific light-detection properties of melanopsin (Lucas et al., 2014), such as its insensitivity and short-wavelength preference, to promote or avoid its activation at different times of day.

However, this approach is problematic because acute activation of melanopsin to promote alertness has the unintended effect of shifting the circadian clock (Provencio et al., 1994), thereby making subsequent sleep difficult. Our identification that the Brn3b(+) ipRGCs specifically mediate light’s acute effects on body temperature provides a cellular basis for developing targeted methods for promoting acute alertness, while minimizing circadian misalignment.

## Methods

### Animals (body temperature)

All procedures were conducted in accordance with NIH guidelines and approved by the Institutional Animal Care and Use Committee of Johns Hopkins University. All mice were maintained on a mixed C57Bl/6J; 129Sv/J background and kept on ad libitum food and water under a 12-hr/12-hr light/dark cycle in group housing until experimentation, with temperature and humidity control. Male and female mice between the ages of 2 and 6 months were used for analysis.

### Body temperature recordings

Each mouse was single-housed at the time of experiment. Surgery was conducted under tribromoethanol (Avertin) anestheshia and a telemetric probe (Starr G2 E-Mitter) was implanted in the peritoneal cavity to monitor core body temperature and general activity. Data was collected in continuous 1- or 2-min bins using VitalVIEW software and analyzed in Microsoft Excel. All experiments were conducted at least 10 days after surgery. Lights were controlled by a programmable timer and all lights were 6500K CFL bulbs illuminated each cage at ~500 lux. Light intensity (Figure S1) was modulated using

*Brn3b*^*Cre*/+^ mice were anesthetized with tribromoethanol (Avertin) and 0.5-1 μl AAV2-hSyn-DIO-hM3Dq-mCherry (UNC Vector Core) was injected intravitreally in one eye using a picospritzer and pulled pipet. At least one week later, animals underwent surgery for implantation of telemetric probes (as above). All experiments were conducted at least 10 days after telemetric probe implantation and at least three weeks after viral injection. After behavior, the eyes of each animal were inspected to ensure that >50% infection had been achieved. We routinely saw >80% of the retinas were infected as we have described previously (Keenan et al. 2016).

Diurnal amplitude was measured by subtracting the mean body temperature for the light cycle (ZT0-12) from the mean body temperature for the dark cycle (ZT12-24). Mean body temperature during testing used all data from ZT14-17. For CNO experiments, injections were carried out near ZT14, but specific times were recorded for each mouse to align the data to the time of injection. Comparisons of mean body temperature after PBS or CNO utilized the 6 hours following injection.

Clozapine-N-oxide (Sigma) was prepared as a 0.1 mg/ml solution in PBS and injected at 1 mg/kg intraperitoneally at ZT14. After behavior, the eyes of each animal were inspected to ensure that >50% infection had been achieved. We routinely saw >80% of the retinas were infected as we have described previously (Keenan et al. 2016).

### Animals (Sleep)

All procedures were conducted in accordance with NIH guidelines and approved by the Institutional Animal Care and Use Committee of Northwestern University. Opn4Cre and Brn3bz-dta were maintained on a mixed C57Bl/6J; 129Sv/J background (Hattar et al., 2002, 2006; Mu et al., 2005). Male and female littermate *Opn4*^*Cre*/+^ and *Opn4*^*Cre*/+^; *Brn3b*^*z-dta*/+^ animals between the ages of 2 and 3 months were used for sleep analysis.

### Sleep recording

Male and female littermate *Opn4*^*Cre*/+^ and *Opn4*^*Cre*/+^; *Brn3b*^*z-dta*/+^ mice were used for sleep recordings. Electroencephalogram (EEG) and electromyogram (EMG) electrode implantation was performed simultaneously at 8 weeks of age. Mice were anesthetized with a ketamine/xylazine (98 and10 mg/kg respectively) and a 2-channel EEG and 1-channel EMG implant (Pinnacle Technology) was affixed to the skull. Mice were transferred to the sleep-recording cage 6 days after surgery, tethered with a preamplifier, and allowed 3 days to acclimate to the new cage and tether. Mice were housed in 12:12 light/dark conditions before and after EEG implantation. EEG and EMG recording began simultaneously at the end of the habituation period, which were displayed on a monitor and stored in a computer for analysis of sleep states. The high pass filter setting for both EEG channels was set at 0.5 Hz and low pass filtering was set at 100 Hz. EMG signals were high pass filtered at 10 Hz and subjected to a 100 Hz low pass cutoff. EEG and EMG recordings were collected in PAL 8200 sleep recording software (Pinnacle Technology) and scored, using a previously described, multiple classifier, automatic sleep scoring system, into 10-sec epochs as wakefulness, NREM sleep, or REM sleep on the basis of rodent sleep criteria (Gao et al., 2016). Light source for all sleep experiments was a 3000 Kelvin light source at 500 lux.

### Wheel-running activity and Masking experiment

Mice were placed in cages with a 4.5-inch running wheel, and their activity was monitored with VitalView software (MiniMitter). Analyses of wheel running activity were calculated with ClockLab (Actimetrics). We used 500 lux light intensity. Mice were initially placed under 12:12 LD masking experiments. Mice were exposed, in their home cage, to a timer-controlled 3-hour light pulse at ZT14-ZT17. Percent activity for each mouse was normalized to its own activity at ZT13 (1 hour before light pulse), and analyzed in 30-minute bins.

### Statistics

All statistical tests were performed in Graphpad Prism or R 3.4.4. Specific tests are listed in the text and figure legends.

---

**Figure S1:**
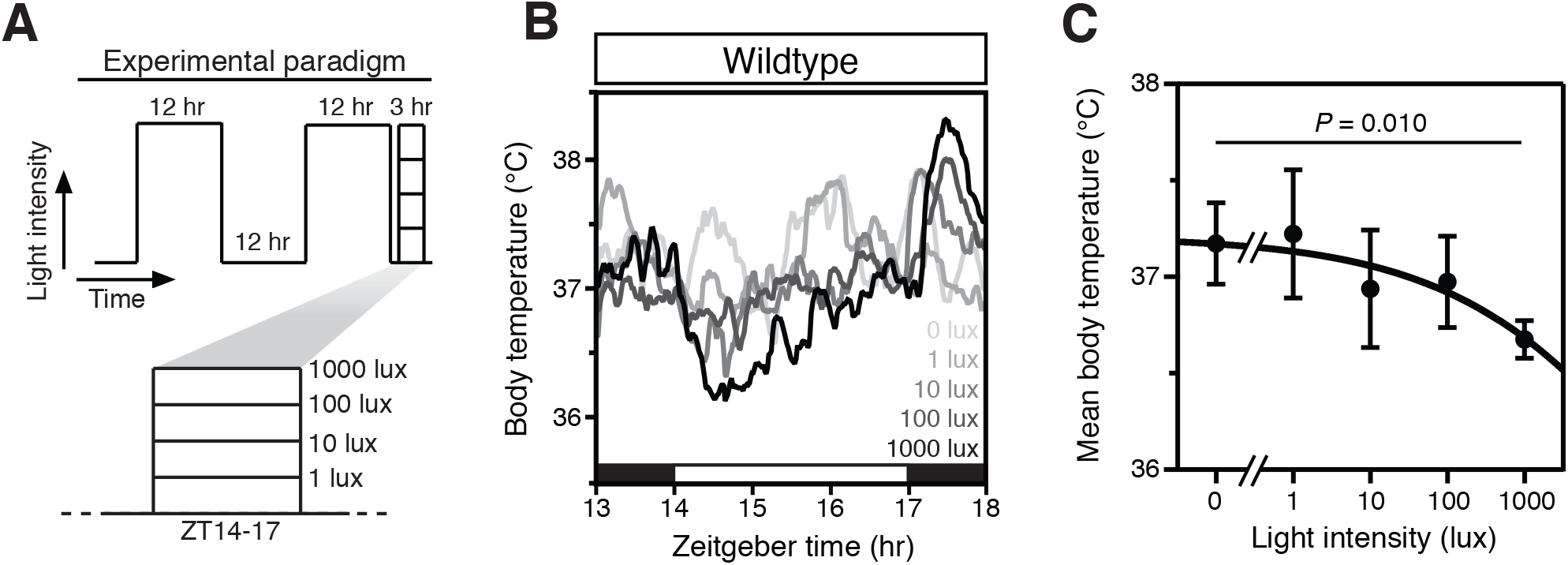
Intensity-dependent decrease in core body temperature during a nocturnal light pulse. (**a**) Experimental paradigm consisting of a 12-hr/12-hr light/dark cycle with a single 3-hr light pulse starting at Zeitgeber time (ZT) 14 (i.e. 2 hours after lights-off). Each experimental night, a light pulse was given at a specific envirnomental light intensity ranging from 1 to 1000 lux in log_10_ increments. (**b**) Mean body temperature for wildtype mice (*n* = 4) that were administered a light pulse at ZT14 of varying intensity (shown as shades of gray). Robust thermoregulation by light only occurs at bright intensities. Black and white bars on the *x* axis refer to time of lights-off and lights-on. (**c**) Quantification of the mean body temperature during the 3-hr light pulse for the wildtype mice in b (*n* = 4, mean ± SD) fit with a sigmoidal dose-response curve. There is a statistically significant effect of light intensity on body temperature (*P* = 0.010), as determined by the main effect of a repeated measures one-way ANOVA.

**Figure S2:**
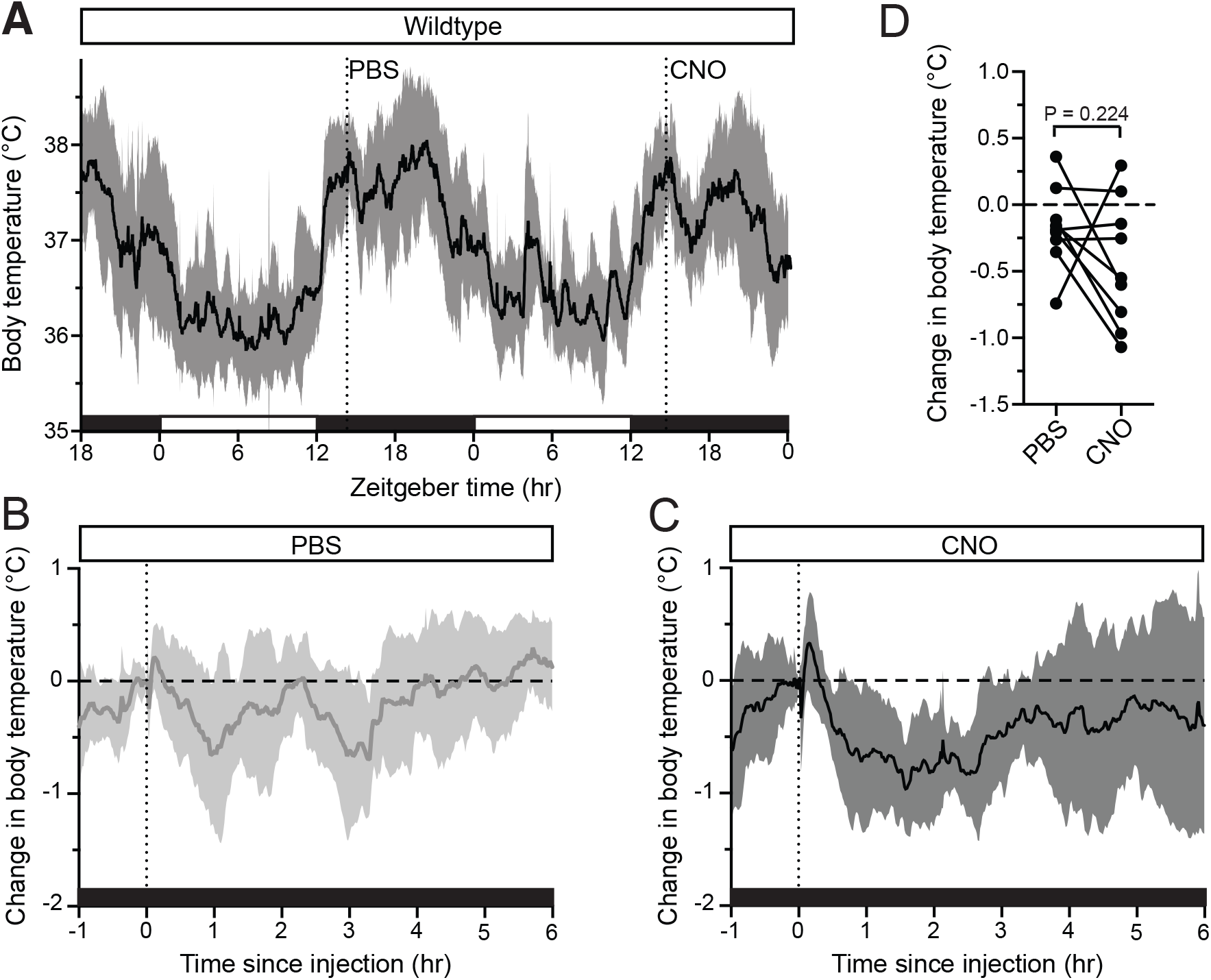
No effect of CNO on body temperature in wildtype mice. (**A**) Wildtype mice body temperature (*n* = 9) was monitored continuously and PBS was injected on night 1 at ZT14, followed by CNO injection (1 mg/kg) on night 2 at ZT14. (**B,C**) Normalized body temperature of either (**B**) PBS or (**C**) CNO injection. Both injections generate a rapid body temperature increase, followed by a dip below the reference value, before returning to normal. (**D**) Paired comparisons of body temperature changes in response to either PBS or CNO administration.

**Figure S3:**
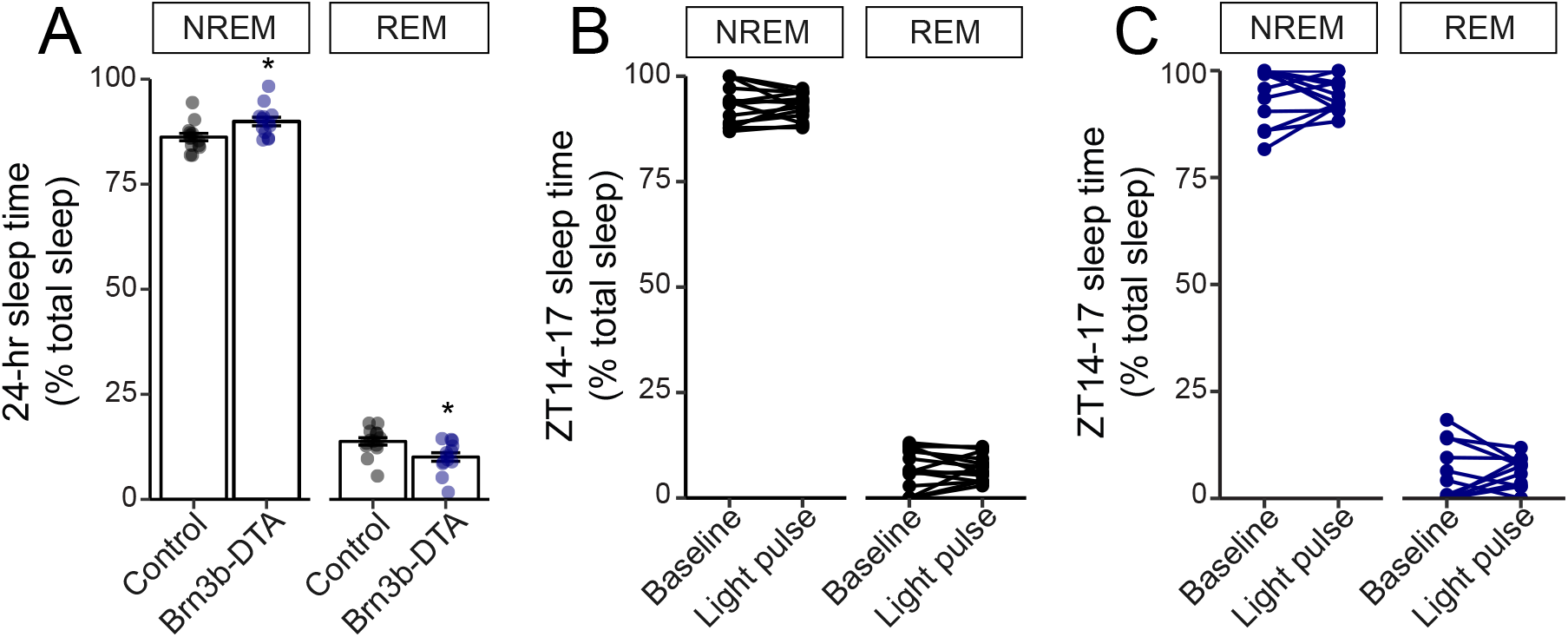
NREM and REM measurements in Control and Brn3b-DTA mice. (A) Percent sleep recorded across the 24 hour day as NREM vs. REM in Control (black) and Brn3b-DTA mice (blue). Brn3b-DTA mice showed a small but significant increase in NREM and decrease in REM sleep compared to Control mice. REM: rapid eye movement, NREM: non-REM. Control: n = 14. Brn3b-DTA: n = 13. \**P* = 0.011 by t-test. (B,C) Sleep stage quantification during ZT14-17, either baseline night or light pulse. No significant differences were seen in either (B) Control or (C) Brn3b-DTA mice by paired t-test.

**Supplemental Figure S4:**
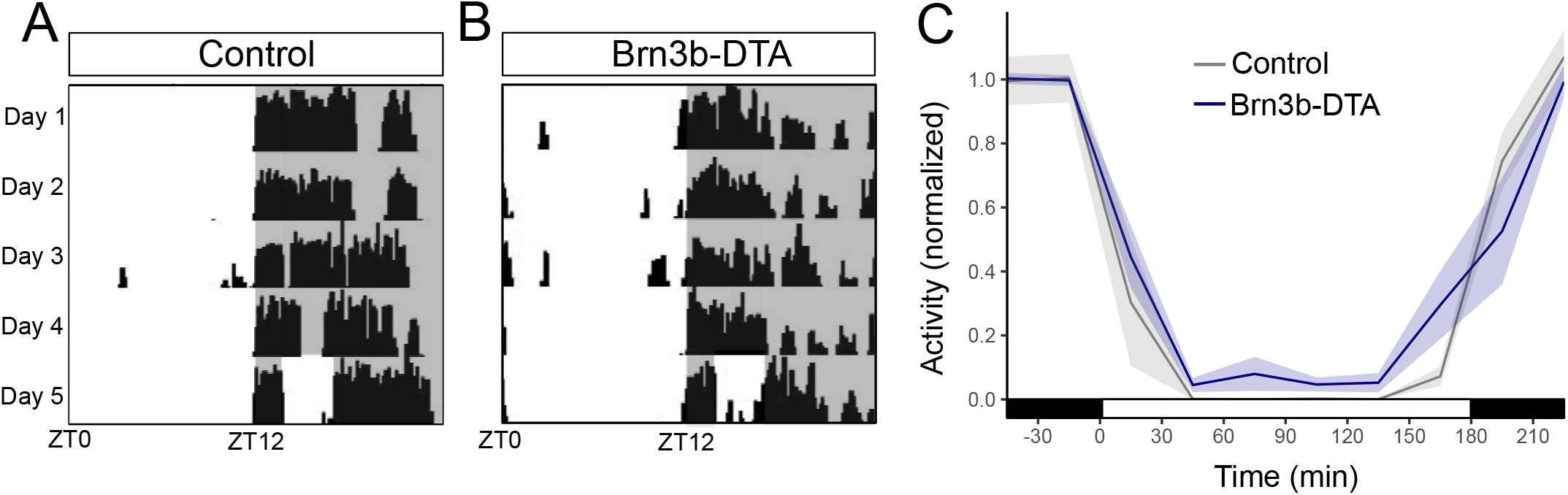
Wheel-running activity in Brn3b-DTA mice. (**A**) Control (Opn4^Cre/+^, n = 5) and (**B**) Brn3b-DTA (Opn4^Cre/+^; Brn3b^DTA/+^, n = 6) were housed in a 12:12 LD cycle and subjected to a 3-hr light pulse starting at ZT14. (**C**) Activity counts in 30 minute bins for both groups. Wheel revolutions were normalized to the average activity for the 1 hour preceding the light pulse. Shading represents SEM. While both groups display robust wheel-running inhibition in response to the light pulse, Brn3b-DTA mice had a mild deficit compared to Controls (*P* = 0.039 by linear mixed model).

## Acknowledgements

This work was supported by a Klingenstein-Simons Fellowship in the Neurosciences, a Sloan Research Fellowship in Neuroscience, and NIH 1DP2EY027983 to T.M.S. and NIH GM076430 and EY024452 to S.H.

